# Defensive forwards: stress-responsive proteins in cell walls of crop plants

**DOI:** 10.1101/2020.02.15.950535

**Authors:** Liangjie Niu, Wei Wang

## Abstract

As the vital component of plant cell wall, proteins play important roles in stress response through modifying wall structure and involving in wall integrity signaling. However, the potential of cell wall proteins (CWPs) in improvement of crop stress tolerance has probably been underestimated. Recently, we have critically reviewed the predictors, databases and cross-referencing of subcellular locations of possible CWPs in plants (*Briefings in Bioinformatics* 2018;19:1130-1140). In this study, taking maize (*Zea mays*) as an example, we retrieved 1873 entries of probable maize CWPs recorded in UniProtKB. As a result, 863 maize CWPs are curated and classified as 59 kinds of protein families. By referring to GO annotation and gene differential expression in Expression Atlas, we highlight the potential of CWPs as defensive forwards in abiotic and biotic stress responses. In particular, several CWPs are found to play key roles in adaptation to many stresses. String analysis also reveals possibly strong interactions among many CWPs, especially those stress-responsive enzymes. The results allow us to narrow down the list of CWPs to a few specific proteins that could be candidates to enhance maize resistance.

## 1. Introduction

Crop plants, especially cereals such as wheat, rice and maize, are the main source for food and feed worldwide. In nature, crops are often suffering from abiotic and biotic stresses, which adversely affect plant growth, development and eventually production. Therefore, developing stress-resistant crops is crucial for global food security [1].

The major abiotic stresses that limit maize production includes drought, heat, cold and flooding [2,3]. Maize yield decreases sharply when the plants are exposed to high temperature (>29°C), particularly in the USA and in northern China [4]. Maize is sensitive to flooding at seedling stage; about 25∼30% annual yield loss is caused by flooding in India alone [5]. It is often plagued by insect pests (e.g., corn borer and nematode) and fungal invasion [6,7]. Alone the fungus *Colletotrichum graminicola*, which induces maize anthracnose, is responsible for annual loss of up to one billion dollars in the USA [8].

In response, plants have developed the sophisticated mechanisms to adapt to both abiotic and biotic stresses, such as forming structural barriers and recruiting chemical compounds. As the outermost layer facing the environments, the cell wall provides stability and protection to the plants and is involved in stress perception [9,10]. The role of plant cell wall in stress response is increasingly emphasized, especially the surveillance of its structure is closely associated with innate immunity in plants [11]. Notably, cell wall proteins (CWPs) have been implied in various stresses through modifying wall structure and involving in wall integrity signaling [12,13]. However, the potential of CWPs in crop improvement of stress tolerance has probably been underestimated.

The composition of cell walls varies with plant species, cell type and developmental stage. Pectin, cellulose and hemicellulose are the main components (>90%) of the primary cell wall, whereas cellulose, hemicellulose and lignin are dominant in the secondary wall [14]. Despite present in minor amounts (5–10% of the primary cell wall mass), CWPs are actively involved in cell wall integrity signaling and innate immunity during plant development and adaptation to environmental cues [12,13]. Most CWPs are basic proteins with a signal peptide and are post-translationally modified, especially by hydroxylation, N-glycosylation and O-glycosylation [14]. After synthesis in the cytosol, CWPs are targeted to the cell wall and/or the extracellular space via the secretory pathways from endoplasmic reticulum (ER) and Golgi apparatus to the cell wall [15].

Identification and cloning of resistance genes is an important prerequisite for targeted breeding of stress-resistance crops, especially through genome editing technologies. To date, only a small number of resistance genes have been identified and modulated in transgenic maize to enhance resistance and improve production [16-21]. All these known resistance genes encode intracellular proteins [22], whereas CWPs have not yet been targeted for crop improvement of stress tolerance. However, many experimental data suggest the role of CWPs in stress response. For example, proteomic analysis in rice and chickpea revealed that many different abundance proteins in extracellular space may be involved in various cellular processes, e.g. cell wall modification, metabolism, signal transduction, cell defense and rescue [23,24].

Over the past 10 years, genome sequencing [25,26] and high-throughput profiling analyses [27-29] in maize have generated huge amounts of CWPs data that have been stored in public databases, especially in the UniProt Knowledgebase (UniProtKB). In this study, we have performed the bioinformatic analysis of CWPs in maize. Our results highlight the potential of CWPs as defensive forwards in stress adaptation. It may provide candidates for targeted improvement of stress resistance in maize and other crops.

## 2. Methods

### 2.1 Retrieving possible maize CWPs entries

UniProtKB is the central hub for the collection of functional information on proteins, with accurate, consistent and rich annotation [30]. Protein entries in UniProtKB have either been confirmed with experimental evidence at protein level or are entirely predicted. Search of UniProtKB with the keyword ‘*Zea mays*+ cell wall or apoplast or secreted protein’ (December 1, 2019) retrieved 1873 entries of possible maize CWPs, with only 50 curated entries. Notably, many sequences were over-represented in UniProtKB, because the fact that different maize lines have been sequenced and submitted separately. Therefore, we have merged redundant sequences and only kept the entries with complete sequences. This applied equally to protein isoforms produced from a single gene.

### 2.2 Curating possible maize CWPs entries

We curated the subcellular locations of all maize CWPs entries retrieved from UniProtKB. Only a small number of the entries are annotated with definite localizations in cell wall and/or extracellular spaces, whereas the majority of the records are computationally analyzed and have no localization annotations. For the latter entries, we predicted their locations with the software HybridGO-Loc [31], as we previously recommended [15]. Only those with definite locations of cell wall or extracellular space were retained for further analysis. For uncharacterized proteins with definite locations of cell wall or extracellular space, BLAST was run against UniProtKB to find their homologues for functional assignment. The gene differential expression of maize CWPs was referred to Expression Atlas (https://www.ebi.ac.uk/gxa/).

### 2.3 Protein-protein interaction analysis

Protein-protein interaction networks were analyzed using the publicly available program STRING (http://string-db.org/). STRING is a database of known and predicted protein-protein interactions.

## 3. Results and Discussion

The cell wall proteome consists mainly of *strict sensu* CWPs present only in the wall and secreted proteins present in the extracellular space (**Fig. 1A**). Inside a plant, the space outside the plasma membrane can be defined as the apoplast where material can diffuse freely. Gene Ontology (GO) definitions of CWPs include cell wall (GO:0005618) protein, apoplast (GO:0048046) protein or secreted protein. Thus, apoplast proteins represent the generalized CWPs. To an extent, cell wall, apoplast and extracellular space are partially overlapping in scope and include related proteins.

**Fig. 1.**
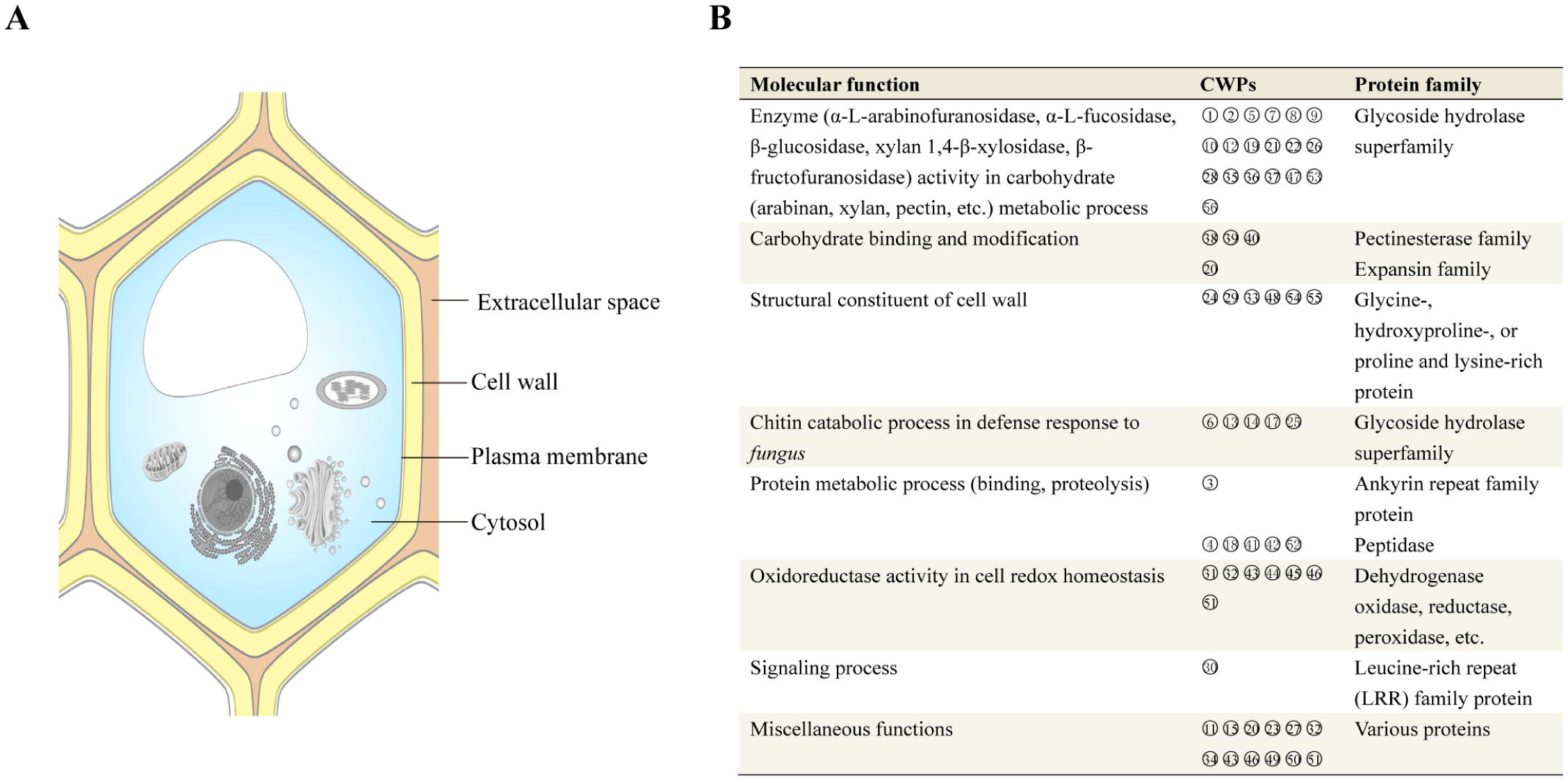
Subcellular localizations highlighting cell wall and extracellular space (**A**) and major molecular functions (**B**) of the examples of maize CWPs. 1, α-L-arabinofuranosidase 1; 2, α-L-fucosidase 2; 3, ankyrin repeat family protein; 4, aspartyl protease AED3; 5, auxin-induced β-glucosidase; 6, basic endochitinase; 7, β-D-xylosidase; 8, β-fructofuranosidase; 9, β-glucosidase; 10, β-hexosaminidase; 11, carbohydrate-binding-like fold; 12, cell wall invertase; 13, chitinase; 14, chitin-binding type-1 domain-containing protein; 15, dirigent protein; 16, DUF1005 family protein; 17, endochitinase; 18, eukaryotic aspartyl protease; 19, exopolygalacturonase; 20, expansin; 21, α-galactosidase; 22, β-galactosidase; 23, germin-like protein; 24, glycine-rich cell wall structural protein; 25, glyco_hydro_19_cat domain-containing protein; 26, glycoside hydrolase; 27, group 3 pollen allergen; 28, heparanase-like protein 3; 29, hydroxyproline-rich glycoprotein (HRGP); 30, leucine-rich repeat (LRR) family protein; 31, malate dehydrogenase; 32, NADH-cytochrome b5 reductase; 33, non-classical arabinogalactan protein 31; 34, nudix hydrolase domain-containing protein; 35, O-glycosyl hydrolase superfamily protein; 36, pectin acetylesterase; 37, pectin lyase; 38, pectin methylesterase 1; 39, pectinesterase; 40; pectinesterase/pectinesterase inhibitor; 41; pepsin A; 42, peptidase A1 domain-containing protein; 43, peroxidase 1; 44, peroxiredoxin; 45, L-ascorbate oxidase; 46, polyamine oxidase 1; 47, polygalacturonase; 48, proline and lysine rich protein; 49, protein EXORDIUM-like 3; 50, purple acid phosphatase (PAP); 51, pyrroline-5-carboxylate reductase; 52, subtilisin-like protease SBT2.6; 53, UDP-arabinopyranose mutase; 54, wall structural protein; 55, vegetative cell wall protein gp1; 56, xyloglucan endotransglucosylase/hydrolase.

After thorough curation, the maize CWPs dataset included 863 entries (Supplementary Table S1), belonging to 56 kinds of protein families (Supplementary Table S2). We functionally classified maize CWPs (**Fig. 1B**) by referring to functional classes of Arabidopsis CWPs [14,32]. According to the number of entries, the top 10 families are expansin (109), pectinesterase (108), xyloglucan endotransglucosylase/hydrolase (XTH) (80), peroxidase (82), polygalacturonase (69), pectin acetylesterase (66), α-L-arabinofuranosidase (64), pectin lyase (51), germin-like proteins (42), and galactosidase (27). Some members of different families are highly similar, such as chitinase (D0EM57) and endochitinase (P29022) (96.44% identity), non-classical arabinogalactan protein (A0A3L6FGF2) and pistil-specific extensin-like protein (B6UHE3) (99.58% identity).

Notably, 36 kinds of the CWPs represent various enzymes, implied in various physiological or biological processes, especially cell wall organization (including polysaccharide biogenesis, degradation and modification), protein proteolysis, cell redox homostasis, and abiotic and biotic stress responses. Moreover, STRING analysis implied that maize CWPs participate in MAPK and Wnt signaling pathways. It is suggested that MAPK and Wnt signaling pathways may play pivotal roles in linking perception of external stimuli with changes in cellular organization or gene expression [33].

The importance of the CWPs that function in cell wall organization is obvious because polysaccharides constitute the largest components of plant cell walls and are constantly subjected to remodeling during plant development or in response to environmental cues. Likewise, glycoside hydrolase family proteins are very important due to their activities to hydrolyze chitin (a primary component of fungus cell walls) to confer resistance to fungus. In addition, the importance of several oxidoreductases (e.g. L-ascorbate oxidase, malate dehydrogenase, peroxidase, and polyamine oxidase) was expected in maintaining cell redox homeostasis that may subjected to change under various stresses [34].

Regarding subcellular localizations, 32 kinds of the CWPs exist only in the cell wall, 11 only in the extracellular space, and 13 both in the cell wall and the extracellular space. The CWPs present in the cell wall, such as structural proteins, may interact with other wall components by non-covalent linkages to form insoluble networks [35]. The CWPs present in the extracellular space, especially between the cell plasma membrane and the cuticle in aerial organs or the suberin layer in roots, may confer to the plant surface waterproof qualities and protection against biotic and abiotic stresses [36,37]. In addition, many CWPs, such as peroxidase (A5H8G4), malate dehydrogenase (B4FRJ1), peroxiredoxin (B6T2Y1), purple acid phosphatase (B4FR72), NADH-cytochrome b5 reductase (B6TCK3), carbohydrate-binding-like fold (A0A1D6FHT0) and peroxiredoxin (B6T2Y1), also have intracellular locations.

It should be noted that the 863 entries of maize CWPs collected here are sufficiently representative. By comparison, the WallProtDB, a specialized collection of cell wall proteomic data [38], records only 2,170 protein sequences from 11 different plant species (without maize). In particular, the 270 entries of rice CWPs in the WallProtDB, belonging to 46 kinds of protein families, are homologous with the corresponding maize CWPs. Undoubtedly, the list of maize CWPs entries remains incomplete. For example, pectinesterase, endoglucanase and endoxylanase inhibitors are pathogenesis-related proteins found in cereals and dicots [39], but only maize pectinesterase/pectinesterase inhibitor (A0A1D6KNZ1) are retrieved from UniProtKB. This is possibly due to the difficulty in proteomic analysis of the CWPs and the lack of complete annotation of all sequenced genes. With the technical advance in cell wall isolation, proteomic analysis of maize cell walls will identify more the ‘missing’ CWPs.

To summarise the potential functions of maize CWPs, we checked gene differential expression of maize CWPs in Expression Atlas (https://www.ebi.ac.uk/gxa/). Analysis of the inducible genes suggested that 36 kinds of maize CWPs are implied with a definite role in various biotic and abiotic stresses (**Table 1**, Supplementary Table S3). In particular, at least ten CWPs are simultaneously involved in 5-7 kinds of stresses, including germin-like protein (B4FAV5), UDP-arabinopyranose mutase (P80607), β-fructofuranosidase (P49174), chitinase (D0EM57), peptidase A1 domain-containing protein (B4G1Q7), peroxidase 1 (A5H8G4), β-D-xylosidase (B4F8R5), eukaryotic aspartyl protease (A0A1D6DSN9), NADH-cytochrome b5 reductase (B6TCK3), O-glycosyl hydrolase superfamily protein (K7V329), and subtilisin-like protease SBT2.6 (C0P6H8). Our analysis also revealed that maize may recruit a different set of CWPs (**Table 1**) to respond to abiotic and biotic stresses, although both stress responses may share common CWPs.

**Table 1.**
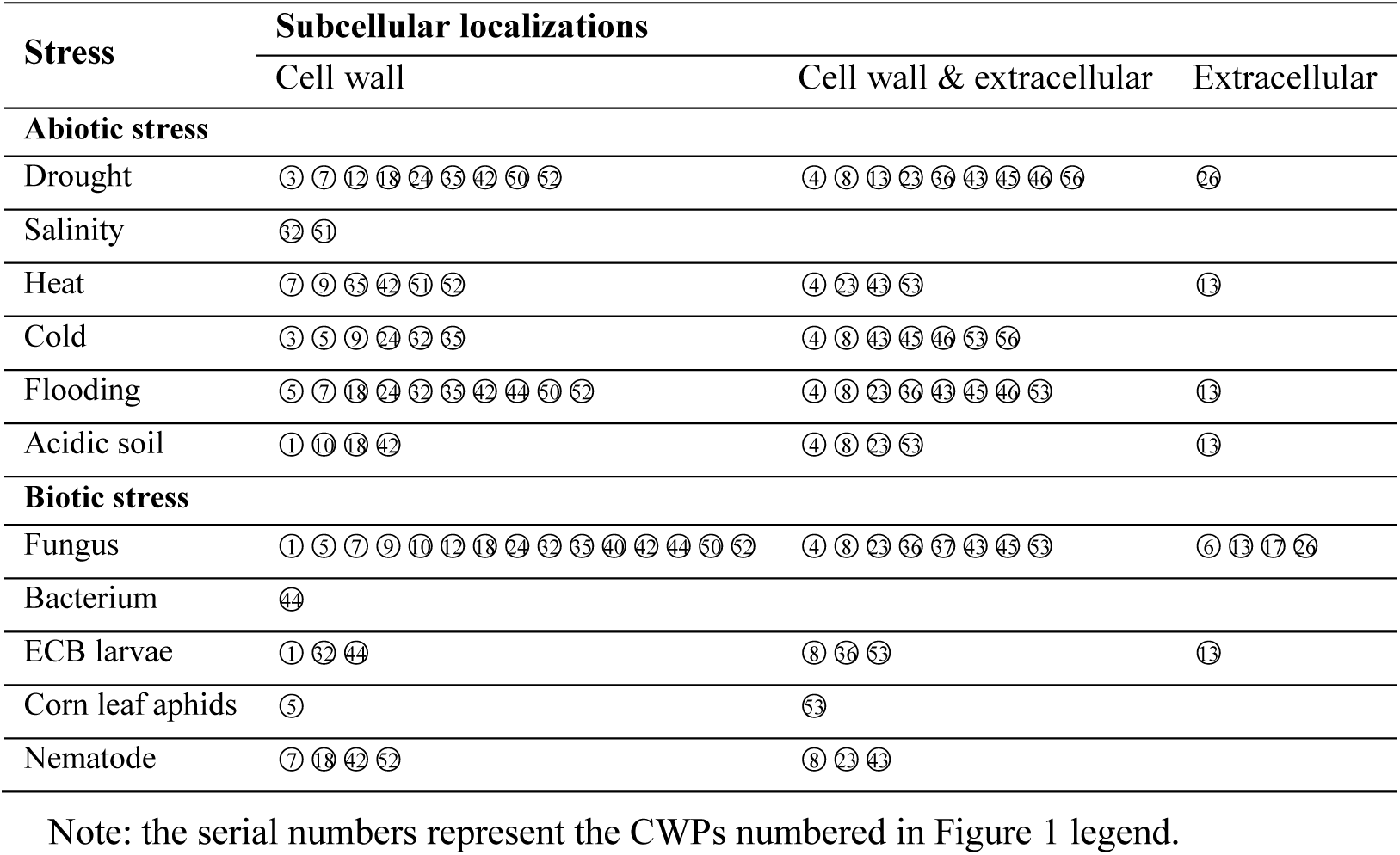
Subcellular localizations of stress-responsive CWPs

The roles of some CWPs in abiotic stresses have been proved in Arabidopsis and other plant species. Here we just referred some examples, because we did not aim to comprehensively review previous studies. In Arabidopsis, pectinesterase 1acts as negative regulators of genes involved in salt stress response [40]; pectin methylesterase is required for guard cell function in response to heat [41]; purple acid phosphatase 17 is reducible by ABA(abscisic acid), H_2_O_2_, senescence, phosphate starvation and salt stress [42]. In rice, β-galactosidase gene responds to ABA and water-stress [43] and germin-like proteins are associated with salt stress [44]. In other plants, β-galactosidase was found to be related to abiotic stress, especially heavy metals [45]; glycine-rich proteins [46,47] and cell wall invertase (copper stress, [48]) are stress-induced. Stress upregulates the expression of expansins and xyloglucan-modifying enzymes that can remodel the wall under abiotic stress [49]. In *Medicago truncatula*, XTH respond to heavy metal mercury, salinity and drought stresses [50], possibly through incorporating newly deposited xyloglucan to strengthen cell walls. However, the role of similar CWPs in maize under abiotic stresses needs to be characterized.

Many maize CWPs are implied to play a role in response of plants to biotic stress. For example, aspartyl protease AED3 (B4FMW6) may be involved in systemic acquired resistance against fungal invasion. Transcription profiling revealed that its transcript (*Zm00001d027965*) was increased by a Log_2_-fold change of 4.3 in maize infected with *Ustilago maydis*. The role of some CWPs in biotic stress have been studied in different plant species. For example, chitinase and endochitinase A have antifungal activity against chitin-containing fungal pathogens [51,52]. Overexpressing extensin enhanced Arabidopsis resistance to *Pseudomonas syringae* by promoting cell wall stiffness [53]. Pectin-degrading enzymes (polygalacturonases, pectatelyases, and pectinmethyl esterases) and xylan-degrading enzymes (endoxylanases) are key virulence factors for pathogens. As a counterattack, plants respond these attacks with a wide range of protein inhibitors of polysaccharide-degrading enzymes [54], e.g. maize pectinesterase/pectinesterase inhibitor (A0A1D6KNZ1) and XIP (xylanase inhibitor protein) [27]. In fruits, polygalacturonases and pectatelyases contribute substantially to the softening of fruit. Suppression of these enzymes delays fruit softening and simultaneously confers enhanced resistance to pathogens like Botrytis [55,56].

As demonstrated in **Table 1**, many maize CWPs may have a role in both abiotic and biotic stress, such as aspartyl protease AED3, HRGPs, β-D-xylosidase, germin-like protein, peroxidase, β-fructofuranosidase etc. It is recognized that HRGPs play major roles in plant defense against abiotic and pathogen attacks [57]. Arabidopsis β-fructofuranosidase (Q43866) is involved in defense to fungus, karrikins and wounding.

However, many maize CWPs did not imply to have significant roles in stress response, such as dirigent protein, expansin, heparanase-like protein, proline rich cell wall protein, exopolygalacturonase, heparanase-like protein 3 etc. However, the roles of dirigent protein and expansins in stress response have been suggested in Arabidopsis [58]. Heparanase activity is strongly implicated in structural remodeling of the extracellular matrix of animals, a process which can lead to invasion by tumor cells [59].

Many protein/protein interactions are expected in cell walls and between CWPs with those spanned in plasma membrane, not to mention leucine-rich repeat (LRR) family protein and pectinesterase/pectinesterase inhibitor enzymes that exist in maize cell walls. Recently, a spanning the plasma membrane protein ZmWAK that confers quantitative resistance to maize head smut, possibly acting as a receptor-like kinase to perceive and transduce extracellular signals [20]. String analysis revealed that many stress-responsive CWPs, especially chitinase, β-hexosaminidase, glycoside hydrolase, α-galactosidase, pectinesterase, and β-fructofuranosidase may form strong-interaction networks in maize (**Fig. 2**). These stress-responsive CWPs can act as a frontline defense or involved in cell signaling process under abiotic and biotic stresses.

**Fig. 2.**
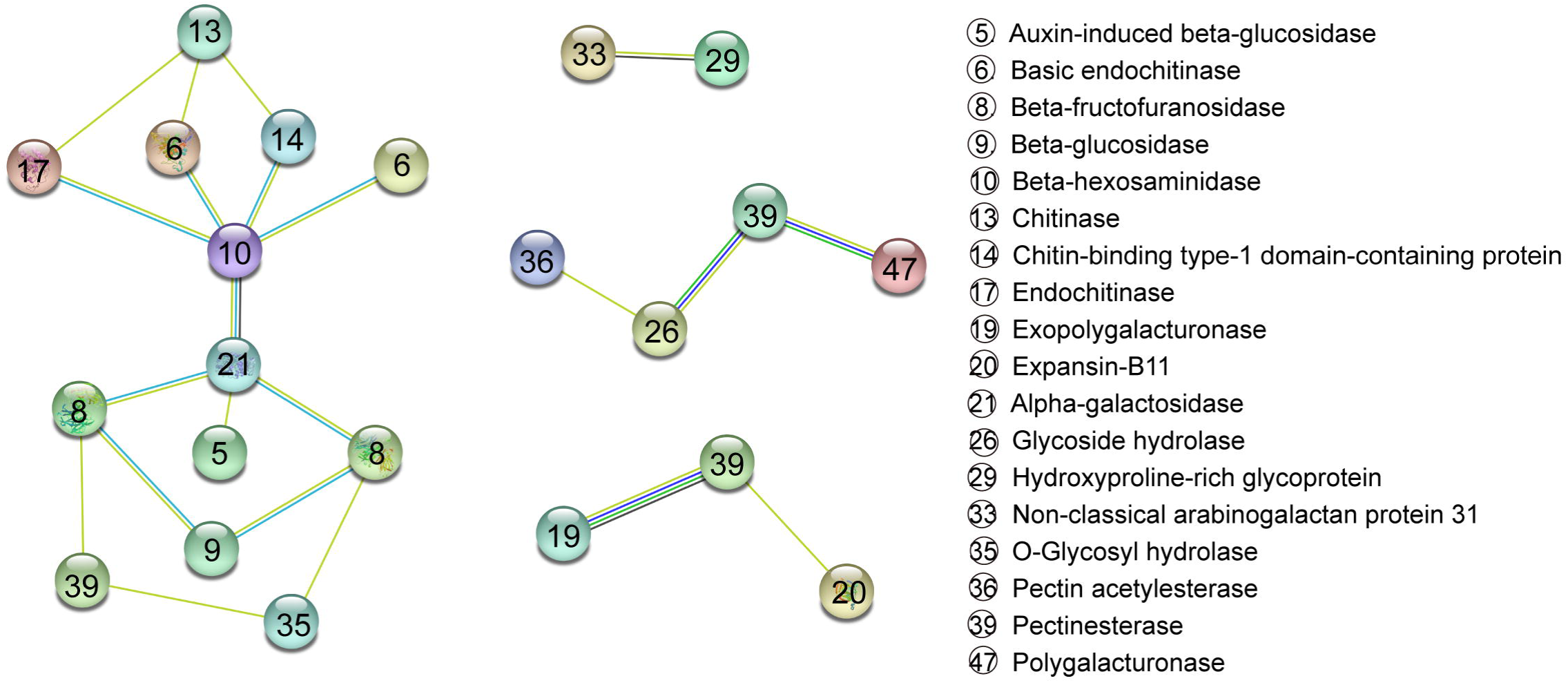
String analysis of possible protein-protein interaction among maize CWPs.

As the interface with the environment, the existing CWPs faces with intense selection pressure to evolve new functions or recruit new proteins to the apoplast through gene duplication and retargeting [60]. Genetic and transgenic evidence in Arabidopsis and other species supported that cell wall gene families associated with cell wall remodeling during abiotic stress and pathogen attack [9]. Therefore, the approach of modifying CWPs provides a novel and rational means of enhancing crop resistance.

At present, many CWPs are suggested as stress-responsive only based on the RNA-Seq data. It was not clear if the protein abundance increased accordingly. So, their accumulation at protein level need to be validated under normal and stress conditions. Another possibility is that not all stress-responsive CWPs play a key role in stress tolerance. This highlights a need to extend genome editing technologies toward specific CWPs. Further molecular and genetic characterization of maize CWPs, especially those with intracellular localizations, will clarify their functions in mediating the plant response to various stresses. Importantly, specific protein abundance in cell walls may not be enough under high stress severity or where crop is exposed to multiple stresses like flooding, drought and diseases, thus enhancing stress-responsive CWPs in crop plants with gene transfer and genome editing is a straightforward approach to enhance crop resistance.

In conclusion, we highlight the potential role of stress-responsive CWPs as defensive forward in maize defense response to various stresses. After clarification of the functions of stress-responsive CWPs during growth, development and stress adaptation, specific CWPs can be candidates for application in genetic modification of stress tolerance in maize and other crops. This may have an important impact on global food security.

## SUPPLEMENTARY MATERIAL

**Supplementary Table S1** Cell wall proteins of maize retrieved in UniProtKB

**Supplementary Table S2** Summary of GO annotation of maize CWPs and their gene differential expression

**Supplementary Table S3** Summary of GO annotation of representative maize CWPs and their gene differential expression

