## Supplemental Table 1-3 for "Defensive forwards: stress-responsive proteins in cell walls of crop plants"

Wei Wang,Weili Huang, Liangjie Niu*

**Supplementary Table S1** Maize CWPs Entries Retrieved in UniProtKB (pages 2-6)

**Supplementary Table S2** Examples of maize CWPs retrieved from UniProtKB (page 7)

**Supplementary Table S3** Summary of GO annotation of representative maize CWPs and their gene differential expression (pages 8-16)

**Supplementary Table S1** Maize CWPs Entries Retrieved in UniProtKB

(Entries are listed in alphabetical order, with reviewed entries in bold and the number in the parentheses represent the number of the sequences)

① Alpha-L-arabinofuranosidase (64): B4FDR3, B4FXI9, B6T9B9, C0P4P5, K7U6T4, K7UM26, A0A1D6F4L8, A0A1D6F4L9, A0A1D6F4M0, A0A1D6F4M2, A0A1D6F4M3, A0A1D6F4M4, A0A1D6F4M5, A0A1D6F4M7, A0A1D6F4M9, A0A1D6F4N0, A0A1D6F4N1, A0A1D6F4N2, A0A1D6F4N4, A0A1D6F4N6, A0A1D6F4N7, A0A1D6FM22, A0A1D6JP53, A0A1D6JP54, A0A1D6K126, A0A1D6K127, A0A1D6K128, A0A1D6K130, A0A1D6K131, A0A1D6K133, A0A1D6KTM6, A0A1D6KTM7, A0A1D6KTM8, A0A1D6NQE9, A0A1D6NQF0, A0A1D6NQF1, A0A1D6NQF3, A0A1D6NQF5, A0A1D6PRI8, A0A1D6QH20, A0A1D6QH21, A0A1D6QH22, A0A1D6QH23, A0A1D6QH24, A0A1D6QH25, A0A1D6QH26, A0A1D6QH28, A0A1D6QH29, A0A1D6QH30, A0A1D6QH31, A0A1D6QH36, A0A1D6QH38, A0A1R3NDP5, A0A317Y633, A0A3L6DAS4, A0A3L6DDF5, A0A3L6DFF5, A0A3L6DSG2, A0A3L6F147, A0A3L6F279, A0A3L6FJ99, A0A3L6FUF4, A0A3L6FYQ6, A0A3L6GC12

② Alpha-L-fucosidase 2 (6): A0A1D6KCJ1, A0A1D6KCJ0, A0A1D6KCI9, A0A1D6KCI8, A0A1D6KCI7, A0A317Y497

③ Ankyrin repeat family protein (4): A0A1D6GGR9, B6TPI5, C0P943, C0P9I4

④ Aspartyl protease/Aspartic proteinase nepenthesin (7): B4FMW6, B4G037, B4G1Q7, B6SJT4, B6TE66, C0P9F8, C0PB10

⑤ Auxin-induced β-glucosidase (1): B6SWK9

⑥ Basic endochitinase (12): B6SZC6, B6T6W1, B6TFQ3, B6TR38, C0P3M6, K7VXP1, A0A1D6GWN1, A0A1D6GWN3, A0A1D6K7T5, A0A1D6LKX7, A0A1D6LMS1, A0A1D6PVW4

⑦ Beta-D-xylosidase (5): B4F8R5, B8A1R0, A0A1D6E4T8, A0A1D6EGK8, A0A1D6JFG2

⑧ Beta-fructofuranosidase, cell wall isozyme (3): **P49174**, A0A1D6H9Z1, A0A3L6EFD5

⑨ Beta-glucosidase (1): A0A1D6GTQ2

⑩ Beta-hexosaminidase (2): B6ST04, A0A1D6LHI9

⑪ Carbohydrate-binding-like fold (1): A0A1D6FHT0

⑫ Cell wall invertase (7): Q9SBI2, Q9SPK0, Q9ZTL2, Q9ZTQ4, Q9ZTQ5, A0A1D6DUP5, A0A1D6J6P6

⑬ Chitinase (3): D0EM57, A0A317Y748, A0A317Y7C2

⑭ Chitin-binding type-1 domain-containing protein (2): B4FK87, A0A1X7YIJ7

⑮ Dirigent protein (1): B6TQI8

⑯ DUF1005 family protein (1): B6SZT8

⑰Endochitinase (7): B4FTS6, B6SZA3, B6TEL0, B6TT00, C0PKN5, **P29022**, A0A1D6E7B3

⑱Eukaryotic aspartyl protease family protein (1): A0A1D6DSN9

⑲ Exopolygalacturonase (18): B4FUB7, B6T4T5, B6TAW9, C4JBB6, **P26216**, **P35338**, **P35339**, A0A317Y5Q5, A0A3L6DI3, A0A3L6E932, A0A3L6E950, A0A3L6EDM2, A0A3L6EHF9, A0A3L6ENL7, A0A3L6EUF7, A0A3L6F4K1, A0A3L6F7R4, A0A3L6FM61

⑳Expansin (109): B4F8B6, B4FEH6, B4FLU4, B4FPA9, B4FRP2, B4FWR8, B4FYF7, B6SGJ2, B6T5Z7, B6T7X4, B6TJY0, B6TP92, B6TQR2, B6UA99, B8A1G1, C0HEC1, C0HEY5, C0P2N8, C0P890, C0P9D0, C0PJW8, C4JBZ0, K7VDB7, K7VT40, K7WH77, **P0C1Y5**, **P58738**, **Q07154**, **Q1ZYQ8**, Q94KT3, Q94KT5, Q94KT6, Q94KT7, A0A096TYJ9, A0A1D6H550, A0A1D6H557, A0A1D6H579, A0A1D6H581, A0A1D6HK98, A0A1D6JBP1, A0A1D6JNZ8, A0A1D6JNZ9, A0A1D6JP00, A0A1D6KM59, A0A1D6KM60, A0A1D6KUN7, A0A1D6KUP3, A0A1D6KUP4, A0A1D6KUP5, A0A1D6KUP6, A0A1D6KUP7, A0A1D6LEA6, A0A1D6LEH7, A0A1D6LEH8, A0A1D6LEH9, A0A1D6LEI0, A0A1D6LEI2, A0A1D6LEI3, A0A1D6LEI4, A0A1D6LEI5, A0A1D6LLK0, A0A1D6M6L1, A0A1D6MPF4, A0A1D6MPF6, A0A1D6MQG9, A0A1D6QEM7, A0A317Y0P1, A0A317Y0P9, A0A317Y1K1, A0A317Y1X3, A0A317Y204, A0A317Y390, A0A317Y3L4, A0A317Y881, A0A317YB29, A0A317YBV3, A0A317YDN7, A0A317YHC7, A0A317YJE0, A0A3L6D8P5, A0A3L6DA75, A0A3L6DEK2, A0A3L6DF32, A0A3L6DGY1, A0A3L6E6H6, A0A3L6E7R1, A0A3L6E7W5, A0A3L6E882, A0A3L6EER6, A0A3L6EGY3, A0A3L6EHG3, A0A3L6EHM1, A0A3L6EKA7, A0A3L6EMW2, A0A3L6ESB1, A0A3L6EVR9, A0A3L6EW90, A0A3L6EWK9, A0A3L6EY84, A0A3L6F4H0, A0A3L6F5I5, A0A3L6FB36, A0A3L6FCH0, A0A3L6FFL8, A0A3L6FL84, A0A3L6FUS7, A0A3L6G407, A0A3L6G9X2, A0A3L6GC21

㉑ Galactosidase, *α-type* (11): B4FA27, C0PAL4, A0A1D6EFC8, A0A1D6F7Z2, A0A1D6HYF2, A0A1D6JPI1, A0A1D6JWU1, A0A1D6KS86, A0A1D6KS92, A0A1D6PKF8, A0A1D6QBS8

㉒ Galactosidase, *β-type* (16): B4F9J1, B8A2F0, C0P3T5, K7V4R8, A0A1D6QQE9, A0A1D6EU33, A0A1D6FPT2, A0A1D6IPT8, A0A1D6JPS0, A0A1D6MYX1, A0A1D6N5T1, A0A1D6N5U0, A0A1D6NGI9, A0A1D6NKE5, A0A1D6PQ20, A0A1D6JWI7

㉓Germin-like protein (subfamily 1, 2, 3 and T) (42): B4FAV5, B4FRS8, B4FUT3, B4FY73, B6UEL1, K7TT00, K7TT05, K7TT15, K7TWM7, K7TWN5, K7U0Q4, K7U0Q9, K7UAX3, K7UAX9, K7UAY3, K7UAZ2, K7UDN3, K7URE9, K7URF1, K7URF5, K7URF9, K7W1E7, Q6TM44, A0A1D6DPN9, A0A1D6DZA4, A0A1D6ELU8, A0A1D6H1F2, A0A1D6H9X6, A0A1D6HTL7, A0A1D6J4J1, A0A1D6JQE7, A0A1D6KN97, A0A1D6KN98, A0A1D6L886, A0A1D6M3P4, A0A1D6N137, A0A1D6NEI1, A0A1D6PUN9, A0A1D6PUP6, A0A1D6PUQ1, A0A1D6Q1Q1, A0A1D6QI94

㉔Glycine-rich cell wall structural protein (14): B4FXY6, B6SPN2, B6SR24, B6ST85, B6TEE0, B6TFA1, B6TY69, B6TZT1, B6U1E3, B6U4A6, B6U6I9, B6UGA1, A0A1D6HBT3, A0A1D6KU97

㉕Glyco_hydro_19_cat domain-containing protein (5): C0HEI0, B6SZN3, C0P306, B8A247,

A0A1D6JS63

㉖Glycoside hydrolase (family 28) (2): B6TX01, B6TXJ8

㉗Group 3 pollen allergen (2): K7U2A7, Q7XBA3

㉘ Heparanase-like protein 1 (8): B4FGA1, B6SRR5, B6U6D3, A0A1D6HNK6, A0A1D6KEH1, A0A1D6NEB8, A0A1D6NTX9, A0A1D6QBZ8

㉙Hydroxyproline-rich glycoprotein (HRGP) (2): B4FHE8, Q42366

㉚Leucine-rich repeat (LRR) family protein (1): K7U7Y3

㉛Malate dehydrogenase (1): B4FRJ1

㉜NADH-cytochrome b5 reductase (1): B6TCK3

㉝ Non-classical arabinogalactan protein 31/ Pistil-specific extensin-like (2): A0A3L6FGF2/B6UHE3

㉞Nudix hydrolase domain-containing protein (1): A0A1D6LN55

㉟ O-Glycosyl hydrolase (12): B4FL15, B4FQC1, B4FU19, B6STC4, C0P2R5, C0PI20, K7V329, K7WB31, A0A1D6LN55, A0A1D6LPQ2, A0A1D6PTZ2, A0A1D6QK01

㊱Pectin acetylesterase (66): B4F9N3, B4F9X6, B4FL51, B4FVG6, B4FZC4, B6T178, B6TXR7, B6U063, B6U7Q4, B8A2J2, C0HHW3, C0P5Y1, C0PGF7, C0PGW1, C0PML3, C4J1M5, K7UYB4, K7V2C5, K7VC42, K7VY73, A0A1D6E0M4, A0A1D6E0M5, A0A1D6E0M6, A0A1D6E0M8, A0A1D6FVP1, A0A1D6FVP2, A0A1D6G797, A0A1D6G798, A0A1D6G799, A0A1D6G7A0, A0A1D6HH87, A0A1D6HH88, A0A1D6IKN4, A0A1D6IKN6, A0A1D6IKN7, A0A1D6IKN8, A0A1D6IKN9, A0A1D6IXV8, A0A1D6K689, A0A1D6LVW0, A0A1D6MMW1, A0A1D6MMW2, A0A1D6MS67, A0A1D6MS68, A0A1D6MZZ9, A0A1D6N558, A0A1D6NME3, A0A1D6NMF0, A0A1D6NMF3, A0A1D6P731, A0A1D6P732, A0A317Y4U1, A0A3L6DJ08, A0A3L6DN84, A0A3L6DQP4, A0A3L6E4G9, A0A3L6E5A0, A0A3L6EJM5, A0A3L6EK46, A0A3L6ETB2, A0A3L6F5A7, A0A3L6FC08, A0A3L6FKB6, A0A3L6FKX2, A0A3L6FTD1, A0A3L6G6D8

㊲Pectin lyase (51): B4F828, B4FQ16, B4FX45, B4G0U5, C0P541, C0PDS0, K7VYM8, A0A1D6FHK6, A0A1D6FIZ2, A0A1D6FL95, A0A1D6FZH5, A0A1D6FZH6, A0A1D6H437, A0A1D6H438, A0A1D6H446, A0A1D6H451, A0A1D6H453, A0A1D6H471, A0A1D6HWL6, A0A1D6I580, A0A1D6JM54, A0A1D6KD22, A0A1D6LAC6, A0A1D6LAC7, A0A1D6LAD1, A0A1D6LRB8, A0A1D6LRC6, A0A1D6LRD7, A0A1D6LRF1, A0A1D6LRF3, A0A1D6LRF7, A0A1D6LRF9, A0A1D6LRK1, A0A1D6LRM6, A0A1D6LRN2, A0A1D6LRN5, A0A1D6LRN7, A0A1D6LRN9, A0A1D6LRP3, A0A1D6LRP4, A0A1D6LRP6, A0A1D6LRQ5, A0A1D6LRQ6, A0A1D6LRQ7, A0A1D6LRQ8, A0A1D6LTC8, A0A1D6MK53, A0A1D6PNR8, A0A1D6Q047, A0A1D6QP73, A0A1Q1AZD7

㊳Pectin methylesterase (5): K7W5K6, A0A1D6J7X2, A0A1D6J7X3, A0A1D6J7X8, A0A1D6J7Y2,

㊴Pectinesterase (108): B4F9U3, B4FCJ7, B4FCM1, B4FCY2, B4FI46, B4FKQ3, B4FKZ4, B4FRR6, B6SZL3, B6TMD5, B6TZ43, B6TZE6, B6U6B3, B8A0X6, B8A1C3, B8A2X5, C0HGE0, C0PFP7, C0PMP7, C4IZT1, C4J3B1, C4J543, K7TP23, K7UUE2, O24596, A0A1D6E3U7, A0A1D6E3U8, A0A1D6E3U9, A0A1D6E3V3, A0A1D6E3V6, A0A1D6E967, A0A1D6E971, A0A1D6E973, A0A1D6E975, A0A1D6E976, A0A1D6E979, A0A1D6E980, A0A1D6E981, A0A1D6E982, A0A1D6E985, A0A1D6F4U1, A0A1D6F4U3, A0A1D6FHT9, A0A1D6FU21, A0A1D6FU22, A0A1D6G099, A0A1D6GCB2, A0A1D6GCB3, A0A1D6GCB4, A0A1D6H4M1, A0A1D6HB34, A0A1D6I082, A0A1D6I4Y9, A0A1D6IJ74, A0A1D6IN03, A0A1D6IP48, A0A1D6IYN8, A0A1D6JFI4, A0A1D6JFI5, A0A1D6JFI7, A0A1D6JFI9, A0A1D6JFJ0, A0A1D6JFJ1, A0A1D6JFJ5, A0A1D6JFJ7, A0A1D6JVC6, A0A1D6JVC7, A0A1D6JVC8, A0A1D6JZZ9, A0A1D6KDB9, A0A1D6KE17, A0A1D6KNK0, A0A1D6MAL2, A0A1D6MAL3, A0A1D6MAL4, A0A1D6MAL5, A0A1D6MAL6, A0A1D6MRY9, A0A1D6N7W7, A0A1D6NCW4, A0A1D6NCW6, A0A1D6NP08, A0A1D6PJ96, A0A1D6PN89, A0A1D6PN90, A0A1D6PN91, A0A1D6PN92, A0A1D6PN93, A0A1D6PN94, A0A1D6PN95, A0A1D6PZP0, A0A1Q1BFQ0, A0A317Y1Q1, A0A317Y8F9, A0A317Y8P7, A0A317YA46, A0A317YGS5, A0A317YIS6, A0A3L6DIV2, A0A3L6DMF5, A0A3L6DST7, A0A3L6DSY6, A0A3L6DY29, A0A3L6E441, A0A3L6F9R9, A0A3L6FDM2, A0A3L6FUS0, A0A3L6GCM3

㊵Pectinesterase/pectinesterase inhibitor (4): A0A1D6KNZ1, A0A1D6N5J3, A0A1D6P828, A0A3L6DFM6

㊶Pepsin A (3): B6SJD9, C0PN95, A0A1D6E4P0

㊷Peptidase A1 domain-containing protein (1): B4G1Q7

㊸Peroxidase (82): **A5H8G4**, B4F7T9, B4FBH0, B4FG25, B4FH35, B4FH68, B4FHG3, B4FK72, B4FKG0, B4FNL8, B4FRD6, B4FSW5, B4FU88, B4FVT1, B4FY83, B4FYH1, B4G1C4, B6SIA9, B6SIU4, B6TWB1, C0HFN4, C0HHA6, C0HIT1, C0P4S1, C0PKS1, C4IZA5, D7NLB3, K7TID5, K7TMB0, K7TMC0, K7U151, K7U159, K7UF86, K7UG68, K7UQ82, K7USX8, K7V8K5, K7VDC0, K7VGN1, K7VGN6, K7VQB0, **Q9FEQ8**, A0A1D6E530, A0A1D6E534, A0A1D6F256, A0A1D6F265, A0A1D6FHV4, A0A1D6FP28, A0A1D6FQT9, A0A1D6FUY7, A0A1D6FUY8, A0A1D6H7N5, A0A1D6HQQ8, A0A1D6HX87, A0A1D6I695, A0A1D6IBW1, A0A1D6IKV7, A0A1D6IKV8, A0A1D6IKV9, A0A1D6IKW0, A0A1D6IKW1, A0A1D6IKW2, A0A1D6IKX3, A0A1D6J1L2, A0A1D6JF04, A0A1D6JNY2, A0A1D6K431, A0A1D6K433, A0A1D6K434, A0A1D6K8V8, A0A1D6KQI0, A0A1D6LE55, A0A1D6LFH4, A0A1D6LYW3, A0A1D6MRI4, A0A1D6MRI5, A0A1D6MSC0, A0A1D6MYJ1, A0A1D6N0K1, A0A1D6N0K3, A0A1D6N9N4, A0A1D6NZH6

㊹Peroxiredoxin (1): B6T2Y1

㊺Plant L-ascorbate oxidase (1): A0A1D6P233

㊻Polyamine oxidase 1 (1): **O64411**

㊼Polygalacturonase (69): B4F8V7, B4FAG1, B4FFH7, B4FMP4, B4FWD4, B4FY93, B4FZT4, B4G1N7,

B6T6X1, B6TDS0, B6TMM0, B6TPQ7, B6TUZ2, B6TYD6, B6U0W6, B6U5Y4, B6UB39, B8A0S9, B8A0S9, B8A1W7, C0P4S3, C0P8E0, C0PIY8, C0PKK6, C4J3A3, C4J705, C4JBH3, K7VC52, K7VKI0, K7W8T8, A0A1D6FI10, A0A1D6FL13, A0A1D6G0A5, A0A1D6L8J1, A0A1D6L8J2, A0A1D6L8S3, A0A1D6MBI7, A0A1D6MBI8, A0A1D6MBI9, A0A1D6MSF0, A0A1D6NHT4, A0A1D6QRB3, A0A1D6QRB5, A0A317Y923, A0A317Y983, A0A317YGJ9, A0A317YH27, A0A3L6D5R1, A0A3L6DD25, A0A3L6DH97. A0A3L6DPM5, A0A3L6DQZ3, A0A3L6DSL2, A0A3L6DV79, A0A3L6DX62, A0A3L6DX72, A0A3L6DXF5, A0A3L6ECQ5, A0A3L6EGR8, A0A3L6ER92, A0A3L6ESU7, A0A3L6EVR8, A0A3L6EYK9, A0A3L6F8E8, A0A3L6FE81, A0A3L6FG77, A0A3L6FGT4, A0A3L6FHN6, A0A3L6FMD2

㊽Proline and lysine rich protein (4): K9L7F0, K9L7S1, K9L8D5, Q9ZNY1

㊾Protein EXORDIUM-like 3 (1): K7V7C0

㊿Purple acid phosphatase (2): B4FR72, B6SWS9

51 Pyrroline-5-carboxylate reductase (1): Q4TZJ2

52 Subtilisin-like protease SBT2.6 (1): C0P6H8

53 UDP-arabinopyranose mutase (2): B4FX25, **P80607**

54 Uncharacterized protein (1): B4FUQ3

55 Vegetative cell wall protein (4): B6SY74, B6T5K5, B6U8J0, A0A1D6MR59

56 Xyloglucan endotransglucosylase/hydrolase (80): B4F837, B4F9C6, B4FAV6, B4FBM2, B4FE99, B4FHS5, B4FSS4, B4FTH5, B4FWJ4, B4FWW4, B4G1Z2, B6T2W7, B6T9E1, B6T9H5, B6TDC2, B6TEW5, B6TH17, B6TJ72, B6TK97, B6TM34, B6TQ18, B6TR01, B6TWP1, B6TX02, B6U5W7, B8A0K3, C0HJ15, C0P2K7, C0P6G9, C0P7W9, C0PC72, C0PCM5, C0PL00, E1U818, K7TQ85, K7TW38, K7U6H7, K7U9U0, K7USG7, K7USS3, K7V8Z6, K7VQE7, K7W8V9, Q42446, Q5JZX2, A0A096T7G6, A0A1D6DXK8, A0A1D6E0F4, A0A1D6E0F6, A0A1D6E0F8, A0A1D6GUH2, A0A1D6GUH3, A0A1D6GUL2, A0A1D6H4U7, A0A1D6IZ08, A0A1D6K9K3, A0A1D6PF72, A0A1D6Q7L1, A0A1D6QTW8, A0A1D6QTX0, A0A317Y4A5, A0A317YG22, A0A317YGM9, A0A3L6DB05, A0A3L6DEA6, A0A3L6DFU1, A0A3L6E5M8, A0A3L6EJS3, A0A3L6EZA2, A0A3L6F3T2, A0A3L6F6F2, A0A3L6F7U3, A0A3L6FNT1, A0A3L6FVI4, A0A3L6FWX7, A0A3L6G4F1, A0A3L6G4Z0, A0A3L6GDI3, A0A1D6IYZ9, A0A1D6IZ02

**Supplementary Table S2** Examples of maize CWPs retrieved from UniProtKB

| No. | Protein | Entry | Gene |
|---|---|---|---|
| ① | Alpha-L-arabinofuranosidase 1 | A0A1D6K126 | 103634588 |
| ② | Alpha-L-fucosidase 2 | A0A1D6KCI9 | Zm00001d030471 |
| ③ | Ankyrin repeat family protein | A0A1D6GGR9 | Zm00001d013242 |
| ④ | Aspartyl protease AED3 | B4FMW6 | 100192462 |
| ⑤ | Auxin-induced β-glucosidase | B6SWK9 | Zm00001d048669 |
| ⑥ | Basic endochitinase | B6TR38 | 100284192 |
| ⑦ | Beta-D-xylosidase | B4F8R5 | 100191418 |
| ⑧ | Beta-fructofuranosidase, cell wall isozyme | P49174 | Zm00001d016708 |
| ⑨ | Beta-glucosidase | A0A1D6GTQ2 | 100279909 |
| ⑩ | Beta-hexosaminidase | B6ST04 | Zm00001d035598 |
| ⑪ | Carbohydrate-binding-like fold | A0A1D6FHT0 | 100384763 |
| ⑫ | Cell wall invertase | Q9ZTQ5 | incw3 |
| ⑬ | Chitinase | D0EM57 | chiA |
| ⑭ | Chitin-binding type-1 domain-containing protein | A0A1X7YIJ7 | 103635126 |
| ⑮ | Dirigent protein | B6TQI8 | Zm00001d004753 |
| ⑯ | DUF1005 family protein | B6SZT8 | 100275827 |
| ⑰ | Endochitinase | P29022 | CHIA |
| ⑱ | Eukaryotic aspartyl protease | A0A1D6DSN9 | Zm00001d001771 |
| ⑲ | Exopolygalacturonase | P26216 | PG1 |
| ⑳ | Expansin | B4F8B6 | 100191303 |
| ㉑ | α-Galactosidase | B4FA27 | 100191764 |
| ㉒ | β-Galactosidase | B4F9J1 | 100191631 |
| ㉓ | Germin-like protein | B4FAV5 | Zm00001d008210 |
| ㉔ | Glycine-rich cell wall structural protein | B4FXY6 | 100280774 |
| ㉕ | Glyco_hydro_19_cat domain-containing protein | A0A1D6JS63 | Zm00001d028116 |
| ㉖ | Glycoside hydrolase, family 28 | B6TX01 | 100284735 |
| ㉗ | Group 3 pollen allergen | K7U2A7 | 103653227 |
| ㉘ | Heparanase-like protein 3 | B4FGA1 | Zm00001d037493 |
| ㉙ | Hydroxyproline-rich glycoprotein (HRGP) | Q42366 | HRGP |
| ㉚ | Leucine-rich repeat (LRR) family protein | K7U7Y3 | 103642195 |
| ㉛ | Malate dehydrogenase | B4FRJ1 | 100272900 |
| ㉜ | NADH-cytochrome b5 reductase | B6TCK3 | 103649684 |
| ㉝ | Non-classical arabinogalactan protein 31 | A0A3L6FGF2 | AGP31_2 |
| ㉞ | Nudix hydrolase domain-containing protein | A0A1D6LN55 | Zm00001d036430 |
| ㉟ | O-Glycosyl hydrolase superfamily protein | K7V329 | 100502339 |
| ㊱ | Pectin acetylesterase | B4F9X6 | 100274042 |
| ㊲ | Pectin lyase | B4F828 | 100191230 |
| ㊳ | Pectin methylesterase 1 | A0A1D6J7X2 | 542462 |
| ㊴ | Pectinesterase | A0A3L6FKB6 | PAE3_1 |
| ㊵ | Pectinesterase/pectinesterase inhibitor | A0A1D6KNZ1 | Zm00001d032180 |
| ㊶ | Pepsin A | A0A1D6E4P0 | Zm00001d002833 |
| ㊷ | Peptidase A1 domain-containing protein | B4G1Q7 | Zm00001d001771 |
| ㊸ | Peroxidase 1 | A5H8G4 | PER1 |
| ㊹ | Peroxiredoxin | B6T2Y1 | Zm00001d046682 |
| ㊺ | Plant L-ascorbate oxidase | A0A1D6P233 | 100273169 |
| ㊻ | Polyamine oxidase 1 | O64411 | MPAO1 |
| ㊼ | Polygalacturonase | A0A3L6DD25 | PGLR_4 |
| ㊽ | Proline and lysine rich protein | K9L7F0 | plr354a |
| ㊾ | Protein EXORDIUM-like 3 | K7V7C0 | 103638128 |
| ㊿ | Purple acid phosphatase (PAP) | B4FR72 | Zm00001d046593 |
| 51 | Pyrroline-5-carboxylate reductase | Q4TZJ2 | p5cr |
| 52 | Subtilisin-like protease SBT2.6 | C0P6H8 | 103629592 |
| 53 | UDP-arabinopyranose mutase | P80607 | UPTG |
| 54 | Wall structural protein | B4FUQ3 | N/A |
| 55 | Vegetative cell wall protein gp1 | A0A1D6MR59 | 103650048 |
| 56 | Xyloglucan endotransglucosylase/hydrolase | B6T2W7 | 100282047 |

**Supplementary Table S3** Summary of GO annotation of representative maize CWPs and their gene differential expression (in Expression Atlas, https://wwwebiacuk/gxa/)

① Alpha-L-arabinofuranosidase 1: A0A1D6K126

GO - Molecular function: α-L-arabinofuranosidase activity

GO - Biological process: L-arabinose metabolic process

Differential expression (Zm00001d028952): response to acidic soil, fungal invasion (*Sporisorium reilianum, F graminearum, F verticillioides)*, and European corn borer (ECB, *Ostrinia nubilalis*) larvae

(Glycoside hydrolase superfamily)

② Alpha-L-fucosidase 2: A0A1D6KCI9

GO - Molecular function: α-L-fucosidase activity

(Glycosyl hydrolase family 95, glycoside hydrolase superfamily)

③ Ankyrin repeat family protein: A0A1D6GGR9

GO: proteasome complex, cell wall, protein binding

Differential expression (zm00001d013242): response to drought, cold

(Ankyrin repeat family protein)

④ Aspartyl protease AED3: B4FMW6

GO - Molecular function: aspartic-type endopeptidase activity

GO - Biological process: protein catabolic process, proteolysis, regulation of programmed cell death, systemic acquired resistance

Differential expression (zm00001d027965): response to fungal invasion (*Ustilago maydis*), drought, heat, cold, acidic soil, waterlogging

⑤ Auxin-induced β-glucosidase: B6SWK9

GO - Molecular function: α-L-arabinofuranosidase activity, xylan 1,4-β-xylosidase activity

GO - Biological process: arabinan/xylan catabolic process

Differential expression (Zm00001d048669): response to fungalinvasion (*F graminearum)*, corn leaf aphids, cold, high nitrate stimulation, waterlogging

(Putative O-glycosyl hydrolase superfamily protein, glycoside hydrolase superfamily)

⑥ Basic endochitinase: B6TR38

GO - Molecular function: chitin binding, chitinase activity, polysaccharide binding

GO - Biological process: carbohydrate metabolic process, cell wall macromolecule catabolic process, chitin catabolic process, defense response to fungus

(Glycoside hydrolase family 19, glycoside hydrolase superfamily)

⑦ Beta-D-xylosidase: B4F8R5

GO - Molecular function: α-L-arabinofuranosidase activity, xylan 1,4-β-xylosidase activity

GO - Biological process: arabinan/xylan catabolic process

Differential expression (zm00001d018080): response to fungal invasion (*F graminearum, U maydis*), nematode *Meloidogyne incognita*, drought, heat, waterlogging

(Glycoside hydrolase family 3, glycoside hydrolase superfamily)

⑧ Beta-fructofuranosidase, cell wall isozyme: P49174

GO - Molecular function: β-fructofuranosidase activity

GO - Biological process: carbohydrate metabolic process

Differential expression (zm00001d016708): response to attack by European corn borer (ECB, *O nubilalis*) larvae, nematode *M incognita*, fungus (*F verticillioides, U maydis*), drought, cold, acidic soil, waterlogging

(Glycoside hydrolase family 32)

⑨ Beta-glucosidase: A0A1D6GTQ2

GO - Molecular function: β-glucosidase activity

GO - Biological process: carbohydrate metabolic process

Differential expression (zm00001d014489): fungal invasion (*F graminearum*), heat, cold

(Glycoside hydrolase 1 family)

⑩ Beta-hexosaminidase: B6ST04

GO - Molecular function: N-acetyl-β-D-galactosaminidase activity, β-N-acetylhexosaminidase activity

GO - Biological process: carbohydrate metabolic process

Differential expression (zm00001d035598): response to fungus (*U maydis*), acidic soil

(Glycoside hydrolase family 20)

⑪ Carbohydrate-binding-like fold: A0A1D6FHT0

GO - Molecular function: carbohydrate binding

⑫ Cell wall invertase: Q9ZTQ5

GO - Molecular function: β-fructofuranosidase activity

GO - Biological process: carbohydrate metabolic process

Differential expression (zm00001d025355): response to fungal invasion, drought

(Glycoside hydrolase family 32)

⑬ Chitinase: D0EM57

GO - Molecular function: chitin binding, chitinase/endochitinase activity, chitin catabolic process

GO - Biological process: carbohydrate metabolic process, cell wall macromolecule catabolic process, chitin catabolic process

Differential expression (zm00001d003190): response to attack by European corn borer (ECB, *O nubilalis*) larvae, fungal invasion (*F verticillioides, F graminearum, Colletotrichum graminicola, U maydis*), drought, heat, acidic soil, waterlogging

(Carbohydrate-binding module family 18, glycoside hydrolase family 19)

⑭ Chitin-binding type-1 domain-containing protein: A0A1X7YIJ7

GO - Molecular function: chitin binding, chitinase activity

GO - Biological process: carbohydrate metabolic process, cell wall macromolecule catabolic process, chitin catabolic process

⑮ Dirigent protein: B6TQI8

GO - Molecular function: guiding stereospecific synthesis activity, identical protein binding, protein homodimerization activity

GO - Biological process: (-)-pinoresinol biosynthetic process, regulation of catalytic activity

⑯ DUF1005 family protein: B6SZT8

With undetermined function

⑰ Endochitinase: P29022

GO - Molecular function: chitin/polysaccharide binding, chitinase/endochitinase activity

GO - Biological process: amino sugar metabolic process, cell wall macromolecule catabolic process, chitin/polysaccharide catabolic process, defense response to fungus

(CBM18, carbohydrate-binding module family 18; GH19, glycoside hydrolase family 19)

⑱ Eukaryotic aspartyl protease: A0A1D6DSN9

GO - Molecular function: aspartic-type endopeptidase activity

GO - Biological process: protein catabolic process, proteolysis

Differential expression (Zm00001d001771): fungal invasion (*F graminearum, U maydis), with* nematode *M incognita,* drought, acidic soil, waterlogging

(Belongs to the peptidase A1 family)

⑲ Exopolygalacturonase: P26216

GO - Molecular function: galacturan 1,4-α-galacturonidase activity, polygalacturonase activity

GO - Biological process: carbohydrate metabolic process, cell wall organization

(GH28, glycoside hydrolase family 28)

⑳ Expansin: B4F8B6

Function: Causes loosening and extension of plant cell walls by disrupting non-covalent bonding between cellulose microfibrils and matrix glucans

GO - Biological process: plant-type cell wall organization; β-expansin B expressed in pollen is involved in sexual reproduction

(Belongs to the expansin family)

㉑ Alpha-galactosidase: B4FA27

GO - Molecular function: raffinose α-galactosidase activity

GO - Biological process: carbohydrate metabolic process

(Glycoside hydrolase family 27)

㉒ Beta-galactosidase: B4F9J1

GO - Molecular function: β-galactosidase activity

GO - Biological process: carbohydrate metabolic process

(Glycoside hydrolase family 35)

㉓ Germin-like protein: B4FAV5

GO - Molecular function: manganese ion binding, nutrient reservoir activity

GO - Biological process: plasmodesmata-mediated intercellular, regulation of root development

Differential expression (zm00001d008210): nematode *M incognita* in root cells, fungal invasion (*F graminearum*, *U maydis, S reilianum*), drought, heat, acidic soil, waterlogging

(Belongs to the germin family)

㉔ Glycine-rich cell wall structural protein: B4FXY6

Differential expression (zm00001d017032): fungal invasion (*F verticillioides*, *U maydis*), drought, cold, waterlogging

(Glycine rich protein family)

㉕ Glyco_hydro_19_cat domain-containing protein: A0A1D6JS63

GO - Molecular function: chitinase activity

GO - Biological process: cell wall macromolecule catabolic process, chitin catabolic process

㉖ Glycoside hydrolase: B6TX01

GO - Molecular function: lyase activity, polygalacturonase activity, virulence factor (domain)

GO - Biological process: carbohydrate metabolic process, cell wall organization (biogenesis/degradation)

Differential expression (zm00001d027441): fungal invasion (*F graminearum, U maydis*), drought

(GH28, glycoside hydrolase family 28)

㉗ Group 3 pollen allergen: K7U2A7

GO - Biological process: sexual reproduction

㉘ Heparanase-like protein 1: A0A1D6KEH1

GO - Molecular function: β-glucuronidase activity

(glycoside hydrolase family 79)

㉙ Hydroxyproline-rich glycoprotein (HRGP): Q42366

GO - Molecular function: structural constituent of cell wall

㉚ Leucine-rich repeat (LRR) family protein: K7U7Y3

㉛ Malate dehydrogenase: B4FRJ1

GO - Molecular function: malate dehydrogenase activity (oxidoreductase)

GO - Biological process: carbohydrate metabolic process

(Belongs to the LDH/MDH superfamily)

㉜ NADH-cytochrome b5 reductase: B6TCK3

GO - Molecular function: copper ion binding, cytochrome-b5 reductase activity, acting on NAD(P)H

GO - Biological process: response to salt stress

Differential expression (zm00001d039656): European corn borer (ECB, *O nubilalis*) larvae, fungal invasion (*F graminearum*, *U maydis),* waterlogging

(Belongs to the flavoprotein pyridine nucleotide cytochrome reductase family)

㉝ Non-classical arabinogalactan protein 31: A0A3L6FGF2

GO - Molecular function: structural constituent of cell wall

㉞ Nudix hydrolase domain-containing protein: A0A1D6LN55

GO - Molecular function: hydrolase activity (cleaving nucleoside diphosphates)

㉟ O-Glycosyl hydrolase superfamily protein: K7V329

GO - Molecular function: α-L-arabinofuranosidase activity, xylan 1,4-β-xylosidase activity

GO - Biological process: arabinan catabolic process, xylan catabolic process

Differential expression (zm00001d052792): fungal invasion (*F graminearum*), drought, heat, cold, waterlogging

(Glycoside hydrolase superfamily)

㊱ Pectin acetylesterase: B4F9X6

GO - Molecular function: pectin acetylesterase activity

GO - Biological process: cell wall organization (biogenesis/degradation)

Differential expression (zm00001d022258): European corn borer (ECB, *O nubilalis*) larvae, fungal invasion (*F graminearum*, *U maydis),* drought, waterlogging

(Belongs to the pectinacetylesterase family)

㊲ Pectin lyase-like superfamily protein: B4F828

GO - Molecular function: lyase activity, polygalacturonase activity

GO - Biological process: carbohydrate metabolic process, cell wall organization (biogenesis/degradation)

Differential expression (zm00001d009341): fungal invasion

(GH28, glycoside hydrolase family 28)

㊳ Pectin methylesterase: A0A1D6J7X2

GO - Molecular function: aspartyl esterase activity, enzyme inhibitor activity, pectinesterase activity

GO - Biological process: cell wall modification

(Belongs to the pectinmethylesterase family)

㊴ Pectinesterase: A0A3L6FKB6

GO – Molecular function: hydrolase activity

GO – Biological process: cell wall organization (biogenesis/degradation)

(Belongs to the pectinacetylesterase family)

㊵ Pectinesterase/pectinesterase inhibitor: A0A1D6KNZ1

GO - Molecular function: aspartyl esterase activity, pectinesterase activity, pectinesterase inhibitor activity

GO - Biological process: cell wall modification

㊶ Pepsin A: A0A1D6E4P0

GO - Molecular function: aspartic-type endopeptidase activity

GO - Biological process: protein catabolic process, proteolysis

(Belongs to the peptidase A1 family)

㊷ Peptidase A1 domain-containing protein: B4G1Q7

GO - Molecular function: aspartic-type endopeptidase activity

GO - Biological process: protein catabolic process, proteolysis

Differential expression (zm00001d001771): nematode *M incognita* in root cells, fungal invasion (*F graminearum, U maydis*), drought, heat, acidic soil, waterlogging

㊸ Peroxidase: A5H8G4

GO - Molecular function: heme binding, metal ion binding, peroxidase activity

GO - Biological process: hydrogen peroxide catabolic process, response to environmental stresses such as wounding, pathogen attack and oxidative stress

Differential expression (zm00001d040702): nematode *M incognita* in root cells, fungal invasion (*C graminicola, F graminearum, F verticillioides, U maydis*), drought, heat, cold, waterlogging

(Belongs to the peroxidase family Classical plant (class III) peroxidase subfamily)

㊹ Peroxiredoxin: B6T2Y1

GO - Molecular function: peroxidase activity, peroxiredoxin activity

GO - Biological process: cell redox homeostasis, defense response to bacterium, response to oxidative stress

Differential expression (zm00001d046682): European corn borer (ECB, *O nubilalis*) larvae, fungal invasion (*F graminearum, U maydis*), bacterium, waterlogging

(Belongs to the peroxiredoxin family Prx5 subfamily)

㊺ Plant L-ascorbate oxidase: A0A1D6P233

GO - Molecular function: copper ion binding, oxidoreductase activity

Differential expression (zm00001d046330): fungal invasion (*F graminearum, F verticillioides, U maydis*), heat, cold, waterlogging

GO - Biological process: cell redox homeostasis, response to oxidative stress

Belongs to the multicopper oxidase family

㊻ Polyamine oxidase 1: O64411

GO - Molecular function: Oxidoreductase, spermidine/spermine oxidase activity, oxidase

GO - Biological process: polyamine/spermine catabolic process

zm00001d024281: fungal invasion (*F verticillioides, C graminicola,*), drought, cold, waterlogging

Belongs to the flavin monoamine oxidase family

㊼ Polygalacturonase: A0A3L6DD25

GO - Molecular function: polygalacturonase activity

GO - Biological process: carbohydrate metabolic process, cell wall organization

Belongs to the glycosyl hydrolase 28 family

㊽ Proline and lysine rich protein: K9L7F0

GO - Molecular function: structural constituent of cell wall

㊾ Protein EXORDIUM-like 3: K7V7C0

GO - Biological process: response to karrikin

㊿ Purple acid phosphatase (PAP): B4FR72

GO - Molecular function: (Hydrolase) acid phosphatase activity, metal ion binding

GO - Biological process: phosphate ion homeostasis

Differential expression (zm00001d046593): fungal invasion (*U maydis*), drought, waterlogging

(Belongs to the metallophosphoesterase superfamily, purple acid phosphatase family)

⑪ Pyrroline-5-carboxylate reductase: Q4TZJ2

GO - Molecular function: (Oxidoreductase) pyrroline-5-carboxylate reductase activity

GO - Biological process: L-proline biosynthetic process, response to heat and salt stress

Belongs to the pyrroline-5-carboxylate reductase family

⑫ Subtilisin-like protease SBT26: C0P6H8

GO - Molecular function: serine-type endopeptidase activity

Differential expression (zm00001d036483): nematode *M incognita,* fungal invasion *(F graminearum, U maydis), response to* drought, heat, waterlogging

Belongs to the peptidase S8 family

⑬ UDP-arabinopyranose mutase: P80607

GO - Molecular function: (Isomerase) UDP-arabinopyranose mutase activity

GO - Biological process: UDP-L-arabinose metabolic process, cell wall organization, cellulose biosynthesis

Differential expression (zm00001d013751): European corn borer (ECB, *O nubilalis*) larvae, corn leaf aphids (*R maidis*), fungal invasion (*F graminearum, U maydis, C graminicola*), drought, heat, cold, acidic soil, waterlogging

CAZy, GT75 Glycosyltransferase Family 75

⑭ Uncharacterized protein: B4FUQ3

GO - Molecular function: structural constituent of cell wall

⑮ Vegetative cell wall protein gp1: A0A1D6MR59

GO - Molecular function: structural constituent of cell wall

⑯ Xyloglucan endotransglucosylase/hydrolase: B6T2W7

GO - Molecular function: hydrolase activity, hydrolyzing O-glycosyl compounds, xyloglucan:xyloglucosyl transferase activity

GO - Biological process: cell wall organization (biogenesis/degradation), xyloglucan metabolic process

Differential expression (zm00001d029814): drought, cold, waterlogging

(Belongs to the glycosyl hydrolase 16 family)
